# Social Disruption: Sublethal pesticides in pollen lead to *Apis mellifera* queen events and brood loss

**DOI:** 10.1101/2020.10.27.354845

**Authors:** Kirsten S. Traynor, Dennis vanEngelsdorp, Zachary S. Lamas

## Abstract

Eusocial *Apis mellifera* colonies depend on queen longevity and brood viability to survive, as the queen is the sole reproductive individual and the maturing brood replenishes the shorter lived worker bees. Production of many crops rely on both pesticides and bee pollination to improve crop quantity and quality. We looked at the resiliency of queens and their brood after one month of sublethal exposure to field relevant doses of pesticides that mimic exposure during commercial pollination contracts. We exposed full size colonies to pollen contaminated with field-relevant doses of the fungicides (chlorothalonil and propicanizole), insecticides (chlorypyrifos and fenpropathrin) or both, noting a significant reduction in pollen consumption in colonies exposed to fungicides compared to control. While we found no difference in the total amount of pollen collected per colony, a higher proportion of pollen to non-pollen foragers was detected in all pesticide exposed colonies. After ceasing treatments we measured brood development, discovering a significant increase in brood loss and/or cannibalism across all pesticide exposed groups. Sublethal pesticide exposure in general was linked to reduced production of replacement workers and a change in protein acquisition (pollen vs. non-pollen foraging). Fungicide exposure also resulted in increased loss of the reproductive queen.

## Introduction

Pollinators mediate the exchange of pollen between flowers, providing a key ecosystem service by ensuring seed and fruit set in 87.5% of flowering plants (1). Eighty-seven of the leading global food crops depend on animal pollination (2). In large agricultural cultivation, this service is primarily provided by managed honey bees, *Apis mellifera*. Critical pollination services by *Apis m.* colonies is valued at $175 billion worldwide (3) and $17 billion in the US (4). To grow these crops, the United States relies heavily on pesticide protection, applying over 1.1 billion pounds of active ingredients annually (5). Much of current commercial production of crops like almonds, apples, blueberries, carrot seeds, and pumpkin, which are completely dependent on honey bee pollination are also intensively treated with pesticides to protect crop quality and reduce pest and disease damage. US farmers spent over $300 billion on pesticides in 2012 (5). Farmers are reluctant to reduce pesticide use for a complex set of reasons, including fear, crop insurance mandates, risk of lower yields and increased administrative burden (6).

While honey bees can encounter pesticides directly while foraging, they also collect pollen from plants and store it inside the colony. Analysis of stored pollen samples reveal that often multiple pesticide residues are found in the same sample. This is concerning as these products would be consumed by nurse bees who use it as their primary protein source. As many as 31 different products were detected in a single pollen sample, with US pollen samples containing a mean of just over 7 in two different surveys (7, 8). Many of these pesticides have non-additive interactions, making their combined risk to honey bee health difficult to assess (9). The stored pollen is consumed primarily by nurse bees, which convert it into proteinaceous glandular secretions called brood food when fed to worker larvae and royal jelly when fed to queen larvae (10, 11). When functioning as a nurse bee, workers will consume over 100 mg of pollen, mostly during the first 10-12 days of life (12, 13). In a sense, nurse bees function as pesticide filters, so even if highly contaminated food is consumed, only trace amounts of contaminates pass to the brood through provisioned brood food (14).

Pesticides exposure can play a role in poor pollinator health (15–17). In addition to direct toxic effects, sublethal exposures are linked to increased incidence of diseases and parasitism (8, 17–22). Combined with other stress factors such as poor nutrition pesticides can have a synergistic negative effect on individual bee longevity (18, 23–25) which can lead to colony decline (26–30). Commercial beekeepers report pesticide exposure as the second most common reason for colony losses, right after other diseases and parasites (31). Field studies looking to measure the effects of pesticide exposure frequently show individual bees in exposed colonies live shorter lives, but fail to show colony level impacts on survivorship (32–34). This “superorganism resilience”, common to highly eusocial organisms like honey bees, is defined as the ability to survive even after the loss of a large portion of worker bees, provided the reproductive queen is maintained (35).

Queen events, occur when colonies rear a new queen to replace an old (and presumably failing) queen, prepare to swarm, or attempt to replace a queen that died suddenly. Such events occur often in honey bee colonies, and colonies that experience them are at greater risk of dying then colonies that do not (26, 36). US beekeepers consider poor queens a leading cause of colony mortality(31, 37). Higher rates of queen events have been reported to occur in certain agricultural setting, for instance the NJ State Apiarist reports significant queen issues during blueberry pollination (Schuler, personal comm.). As a result of increased queen losses and other poor health outcomes occurring post pollination contract, some commercial beekeepers have stopped pollinating blueberries (38). When colonies go queenless or bloodless, they reduce nectar and pollen foraging (39, 40), potentially negatively impacting the number of floral visits and the resulting fruitset (41, 42).

Here, we sought to understand the impacts of long-term sublethal pesticide exposure had on brood viability and queen events in exposed colonies. We investigated the effects of sublethal pesticide exposure of two of the most commonly found insecticides (the organophosphate chlorpyrifos detected in 75% of samples and the pyrethroid fenpropathrin detected in 18% of samples (7)), and two of the most widely applied fungicides on blueberry crops (chlorothalonil detected in 53% of samples in a nationwide survey and in 73% of pollen samples from blueberry (8) and propicanizole detected in 45% of samples collected during blueberry pollination) (7, 8, 43) (Table 1). According to the U.S. Geological Survey, in 2017 the country applied 11-13 million pounds of the fungicide chlorothalonil for agricultural use with 4.3-4.5 million pounds applied on vegetable and fruit crops like blueberries, 2.1-2.5 million pounds of the fungicide propicanizole, 5.5-9.5 million pounds of the insecticide chlorpyrifos, and 0.12-0.125 million pounds of the insecticide fenpropathrin (44), which provides bees ample opportunity to contact these four compounds. All four have previously been found in bee pollen, appearing frequently in pesticide screens of pollen stores in honey bee colonies (Table 1).

**Table 1.** Actual bee bread contamination rates detected in migratory beekeeping colonies in Traynor et al 2016.

| **Insecticide** | **Class** | **Prevalence** | **Mean LD<sub>50</sub>** | **Max LD<sub>50</sub>** | **Used in Experiment** |
| --- | --- | --- | --- | --- | --- |
| Chlorpyrifos | organophosphate | 34.0% | 4.4% | 30.8% | 5.0% |
| Fenpropathrin | pyrethroid | 19.0% | 4.9% | 34.0% | 5.0% |
| **Fungicide** | **Class** | **Prevalence**<br>**in Blueberries** | **Mean LD<sub>50</sub>**<br>**in Blueberries** | **Max LD<sub>50</sub>**<br>**in Blueberries** | **Used in**<br>**Experiment** |
| Chlorothalonil | fungicide | 72.7% | 0.85% | 2.2% | 1.0% |
| Propiconazole | fungicide | 45.5% | 0.02% | 0.05% | 1.0% |

Prevalence is the percentage of samples that were contaminated with a given pesticide. The mean and max LD<sub>50</sub> % values represent how much of their respective LD<sub>50</sub> bees consume, assuming they eat 100 mg of pollen as a nurse bee. A pesticide LD<sub>50</sub> is met and results in a 50% kill when the pesticide contamination rate in ppb divided by the LD<sub>50</sub> = 10,000, as ppb are measured in ug/kg and LD<sub>50</sub> is measured in ug/mg (1 kg = 1,000,000 mg and 1,000,000 mg/100 mg = 10,000). The EPA includes a safety factor of 1/10th when setting food pesticide tolerances. Thus when ppb/LD<sub>50</sub> = 1,000, a bee will consume 10% of her LD<sub>50</sub> during her 10 d nursing phase and sublethal effects are possible.

To determine the impacts of sublethal ingestion of contaminated pollen on colony health and foraging behavior, we thus exposed colonies to sublethal pesticides in pollen under four different treatment exposures: the two fungicides chlorothalonil and propicanizole (fungicide); the two insecticides chlorpyrifos and fenpropathrin (insecticide); the pesticide cocktail of both fungicides and both insecticides (both), compared to an uncontaminated control (control). We quantified the total pollen collected by the colony using pollen traps, the number of pollen vs. non-pollen foragers returning to colonies, and the amount of the provisioned (both contaminated and not) pollen patties consumed. To understand the long-term effects of pesticide exposure on the next generation of workers post exposure, we investigated the impact on hypopharyngeal glands of bees, dissecting 7-day old nurse bees, followed brood viability, plus measured colony strength and incidence of queen events during and after pesticide exposure.

## Results

### Colony Strength

All 24 colonies survived throughout the experiment from July through October, 2018. Colony strength, as measured by frames of bees and frames of brood (45), were recorded monthly. As expected, the colony strength measures varied over time with colonies shrinking overtime in preparation for winter. These populations trends were not different between treatment group (Fig S1) for frames of bees or frames of brood. However, there was an interaction of treatment and time for frames of bees, as both the control group and the insecticide exposed group had substantially more frames of bees in September than the groups exposed to just the two fungicides as well as the colonies receiving all four pesticides.

### Pollen Consumption & Foraging

To understand differences in protein acquisition and use, we analyzed consumption of provisioned (treated or not) protein patties, colony pollen collection via pollen traps and foraging rates. We analyzed provisioned pollen patty consumption (up to 320g per week and 2,240 g over the entire experiment) using a generalized linear model (GLM) with normal distribution for consumption and date and treatment as factors. Provisioned pollen consumption varied over time, but did not interact with or vary by treatment group (Fig. S2). When pollen patty consumption was pooled data across all dates (Fig. 1), the resulting model revealed that treatment had a significant effect on provisioned pollen consumption, with fungicide only treated colonies consuming significantly less than control colonies (Control = 92.4% ±1.4% SE consumed; Fungicide = 83.2% ± 3.4% SE). Colonies exposed to a cocktail of insecticides and fungicides consumed significantly less than colonies in all other treatment groups (Both = 80.5% ± 1.8% SE. The insecticide fed group did not differ from control or the fungicide group (Insecticide = 88.1 ± 1.4% SE)

**Figure 1.**
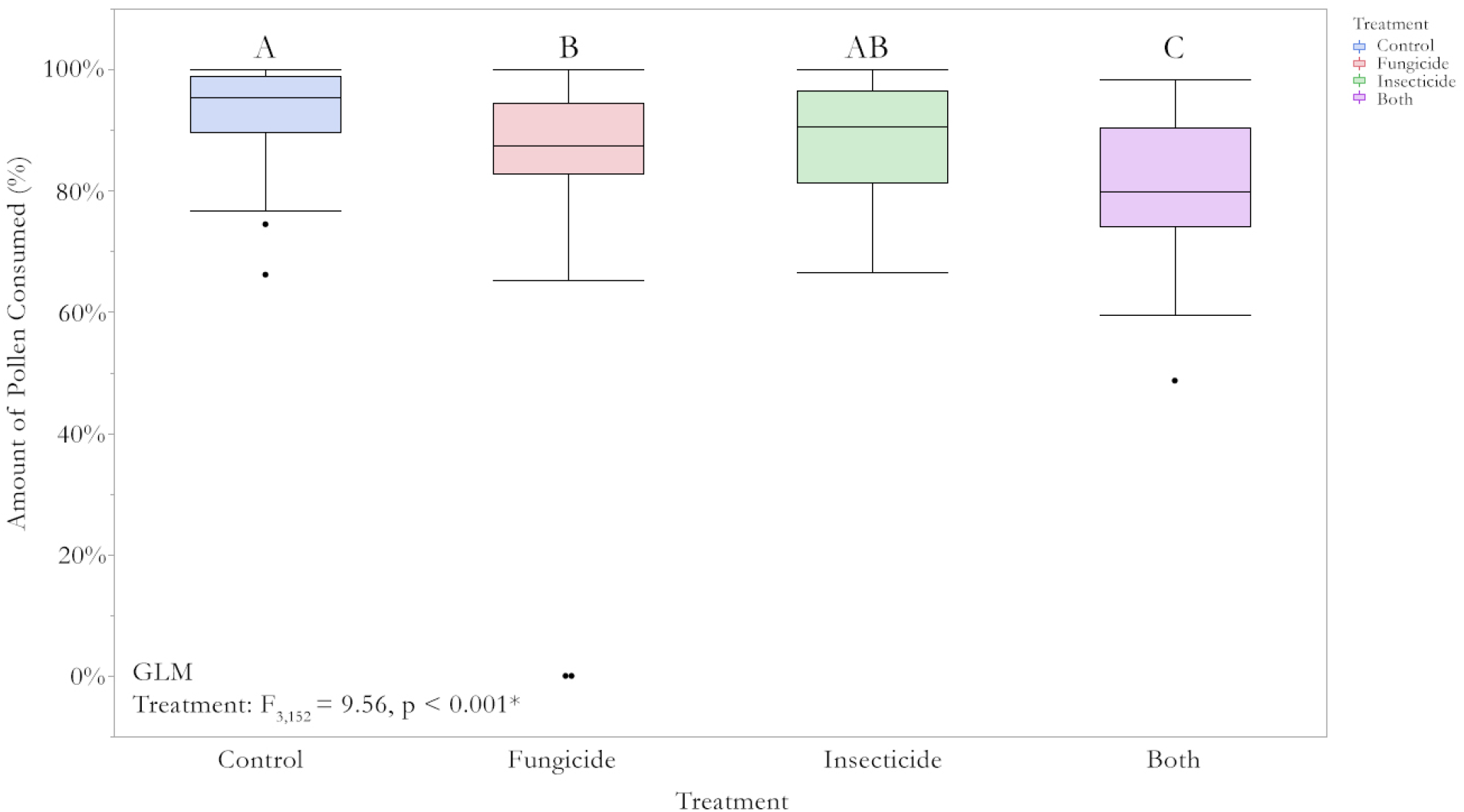
Pollen Consumption over Time. Each colony was provided with four 80g fresh pollen patties 2x per week, typically on the Tues and Saturday, unless it rained. At each indicated date the remainder of the patties was removed and replaced with fresh ones. Colonies could thus consume up to 320g of pollen per feeding. Consumption was analyzed using a GLM with treatment and date as factors. Pollen consumption differed over time (Fig S1), but did not vary significantly by treatment group. Since there was no interaction of treatment and date, we pooled consumption across the entire trial. Consumption than varied significantly by treatment group, with control colonies consuming more than all other groups except the insecticide group. Control = blue, fungicide = red, insecticide = green, both = purple. Box plots show quartiles and outliers. Significant differences indicated by different letters (α = 0.05).

To help ensure consumption of treatment provisioned pollen patties and severely reduce access to foraged pollen, we installed pollen traps on all colonies prior to the start of the experiment. These traps are designed to remove the majority of pollen from the corbicula of returning pollen foragers. We collected and weighed the removed pollen pellets every 3 to 4 days for the 28 days of pesticide treatment, though due to weather we were only able to measure pollen collected on five occasions when the collecting period was uninterrupted by rain. We disengaged the traps so foragers could return with fresh pollen five days after we removed the last treated provisioned pollen patties. We analyzed the amount of pollen collected over the course of the experiment using a GLM with a normal distribution for collected pollen weight and using treatment and date as factors. The amount of pollen collected varied over time, but did not differ by treatment group (Fig. 2).

**Figure 2.**
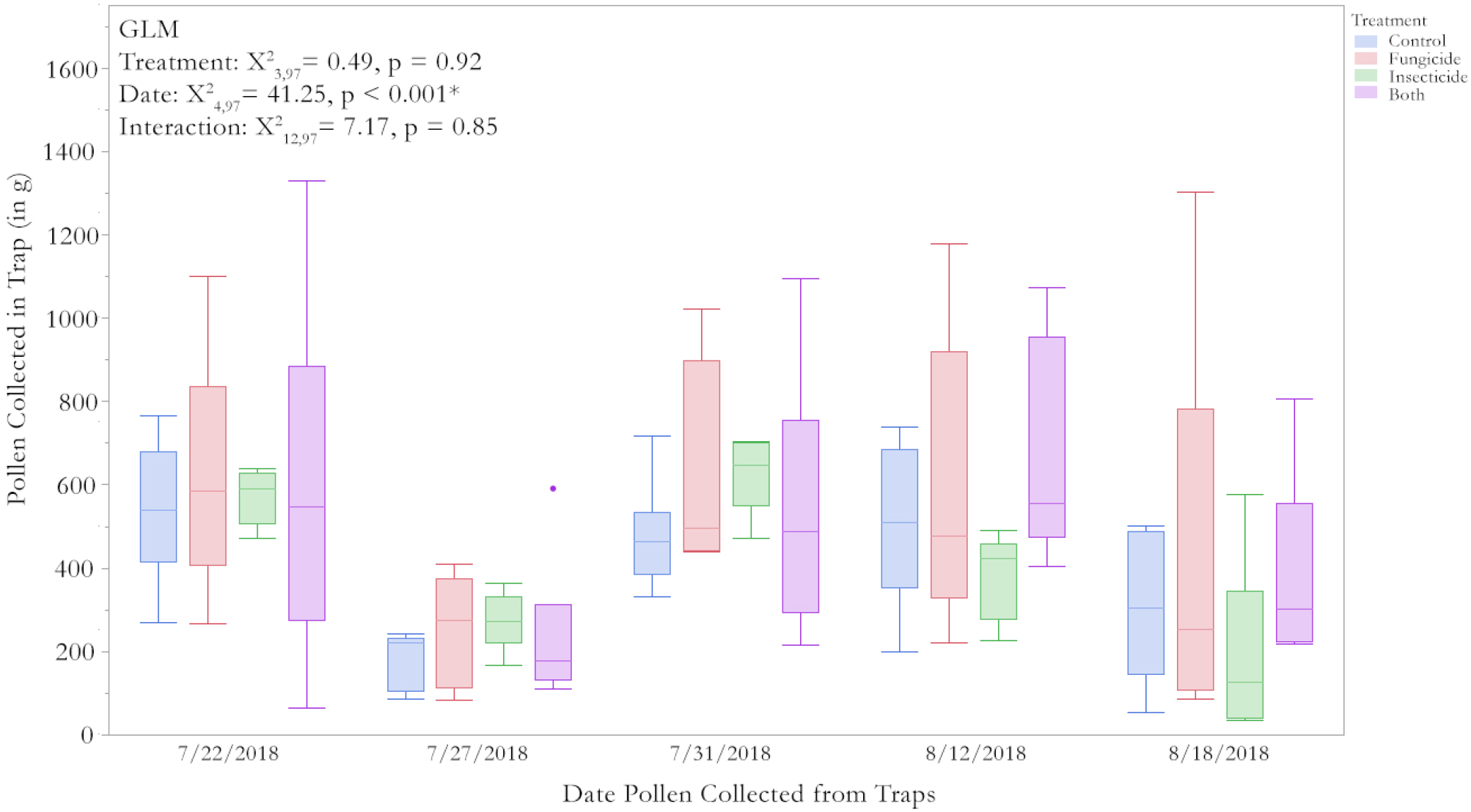
Amount of Pollen trapped per colony. To assure that colonies consumed the pollen patties we fed and did not have access to large amounts of alternative pollen from the environment, each colony was fitted with a Sundance pollen trap that removed the majority of pollen from returning foragers. We trapped pollen from these colonies for 48-72 hour intervals weekly, discarding samples that molded during periods of heavy rain. The amount of pollen collected varied significantly by collection date, but not by treatment group, indicating that the colonies did not differ in the amount of pollen they brought back to the colony. Box plots showing mean, quartiles and outliers, six colonies per treatment group. Pollen traps were permanently removed after our final collection date, as we were no longer feeding the colonies contaminated pollen. Control = blue, fungicide = red, insecticide = green, both = purple.

Colonies exposed to sublethal levels of pesticides could reduce their foraging effort, sending out fewer pollen foragers as a ratio of overall foraging force. To determine this ratio we counted the number of returning foragers at colony entrances for set periods of time, classifying returning bees as either a pollen forager (if they had visible loads of pollen on their corbicula) or non-pollen foragers (if they returned without a pollen load)(39, 46). We analyzed the proportion of returning pollen foragers assuming normal distribution using a GLM with treatment and date as factors (Χ^2^ = 17.82, df = 15, p = 0.27). Since there was no effect of treatment or date and they did not interact, we pooled all data by treatment group to compare the ratio of pollen foragers vs. non-foragers using a Χ^2^ test in a contingency table (Fig. 3). Control colonies had proportionally fewer pollen foragers when compared to each of the pesticide exposed groups, even though there were no differences in the frames of brood, a known key driver of pollen foraging (47).

**Figure 3.**
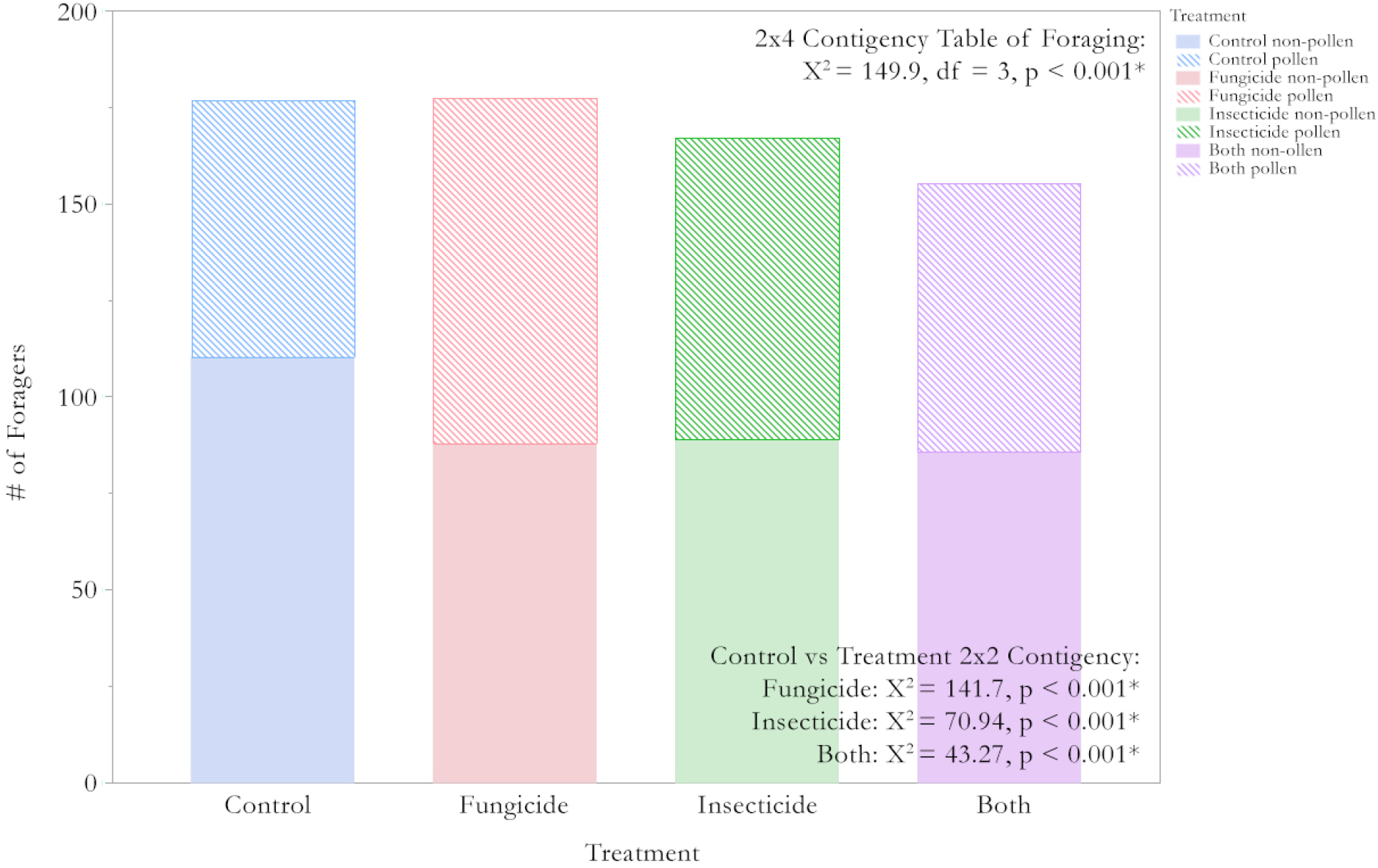
Pollen & Non-Pollen Foraging. We measured the number of pollen foragers and non-pollen foragers returning to each colony during 5-minute intervals on four separate dates. When analyzed with both treatment and date as factors, there was a significant effect of date, but no effect of treatment or interaction. Since treatment and date did not interact, we pooled all data by treatment group to compare the number of pollen foragers vs. non-foragers in a 2×4 contingency table. As that was significant, we then compared the control group to each separate treatment group using a 2×2 contingency table. All of them were significantly different from controls. The control group had a higher proportion of non-pollen foragers compared to each treatment group. Control = blue, fungicide = red, insecticide = green, both = purple. Solid = non-pollen foragers; hatched = pollen foragers.

### Nurse Bees

The colony is dependent on nurse bees to provide protein rich brood food to developing larvae, which is produced predominantly in the hypopharyngeal glands (HPG) of bees. To compare nursing activity among treatment groups we compared HPG size in seven day old bees reared in each of the experimental colonies (48). We noted that some bees (N=2, 1.8% of all bees examined) in the fungicide treated colonies group (8.7% of fungicide bees) had completely atrophied HPG with no glands available for dissection in the head capsule, while other bees from the fungicide group had normal HPG. The number of atrophied glands in fungicide bees (n = 2 of 23) compared to all other groups (n = 0 of 84) via a Fisher’s exact test using a contingency table is significant (p = 0.045). We analyzed HPG differences for all available glands using an ANOVA with colony as a random factor. As we can only compare gland size across bees that had glands available for dissection, we do not find significant differences in HPG size across treatment groups (Fig. 4).

**Figure 4.**
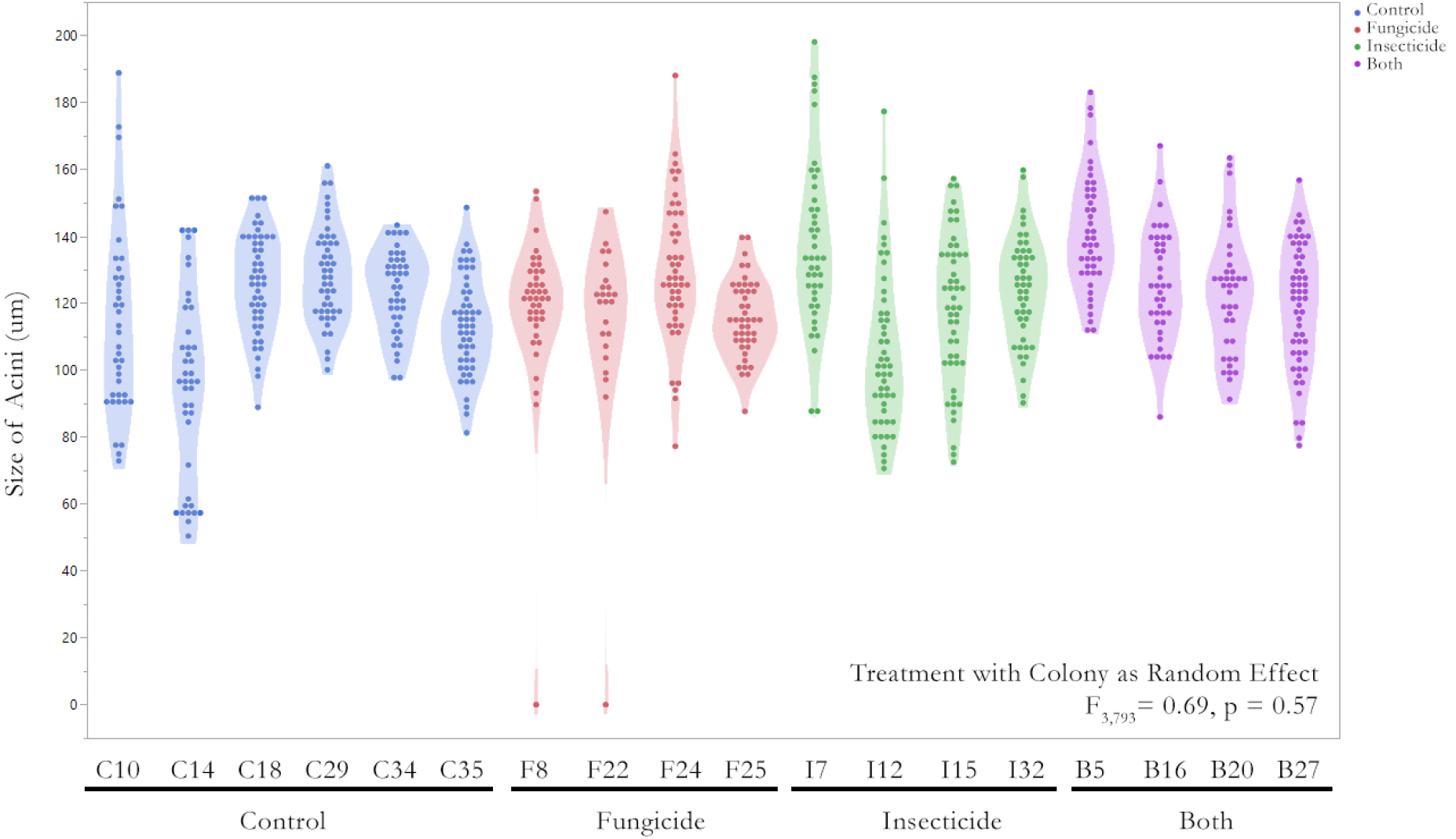
Hypopharyngeal Glands of Nurse Bees. We measured the width of individual acini of 7-day old bees that developed and emerged post pesticide exposure. For each dissected gland, we measured the width of 3-10 acini, depending on how many were in the focal plane. Size differences were compared across treatments with colony as a random effect. There was no significant effect of treatment, though a number of bees in the fungicide group had completely atrophied HPG, making measurement of them impossible. Control = blue, fungicide = red, insecticide = green, both = purple.

### Brood Viability & Queen Events

To measure possible long-term effects of sublethal exposure of pesticides on nurse bees, we monitored the brood viability in colonies after treatment removal, testing 5 days after. This ensured the colony’s nurse bees were reared during treatment application. To measure brood viability, we caged the queens onto an empty brood comb for 24 hours and then monitored the fate of 50 eggs to emergence. Using a repeated measures MANOVA, we found that while time and treatment did not interact, each had a significant effect (Fig. 5a). Colonies in the insecticide treatment group had the highest rates of brood loss, followed by colonies that received the four-pesticide cocktail treatment, and the fungicide only treatment, while control colonies display the least brood loss. We analyzed each stage of development separately, comparing brood loss rates with a normal distribution using a generalized linear model across treatment groups (Fig. 5b and S3). The rate of loss during the larval stage differed between treatment groups, with the insecticide exposed groups having significantly higher rates of loss than the control or fungicide exposed groups. While we did not find any difference in loss rates during the pupal stage, when comparing cumulative loss across all brood stages there was a significant effect of treatment with the insecticide only group having more total brood loss than the control group.

**Figure 5.**
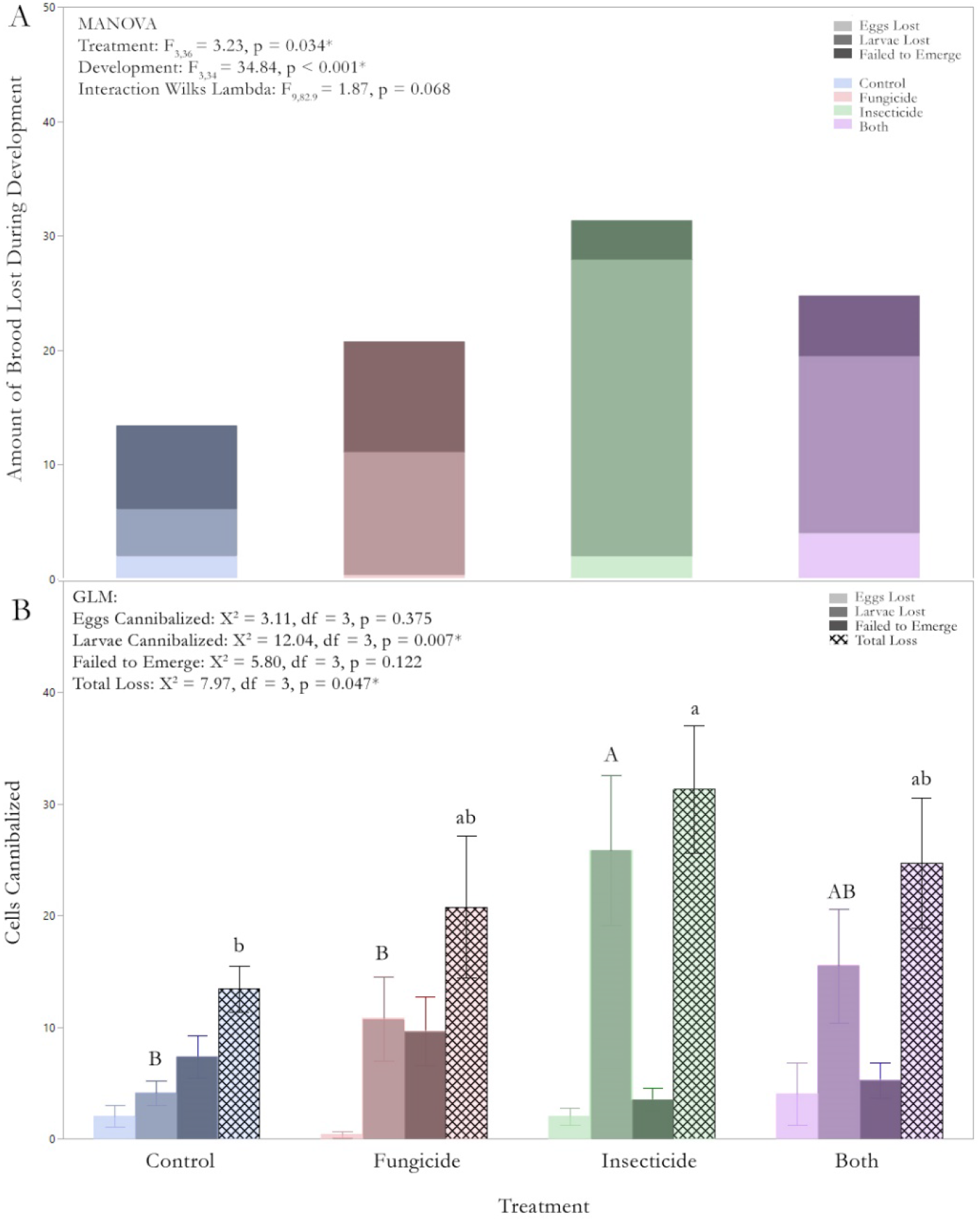
Brood Loss During Development. We caged each queen for egg laying and then followed the development of 50 eggs until emergence. A) We analyzed brood loss over time using a repeated measures ANOVA. Each stacked bar represents the mean brood loss per developmental stage per treatment group over time. Control = blue, fungicide = red, insecticide = green, both = purple. B) Since our repeated measures analysis showed a treatment effect, but no interaction effect, we analyzed the loss at each brood stage individually (egg, larvae, capped brood/failed to emerge) as well as total loss over the entire development period. There was a significant effect of treatment at the larval stage, with significant differences (α = 0.05) between groups represented by capital letters, and for total loss represented by the lowercase letters. Control = blue, fungicide = red, insecticide = green, both = purple. Light bar = eggs lost, medium bar = larvae lost, dark bar = failed to emerge, hatched = total loss.

To quantify queen events, we inspected colonies monthly from July through October. We analyzed the number of queen events using a generalized linear model with a binomial distribution, using month and treatment as factors (Fig 6). Treatment had a significant effect on the number of queen events, but queen events did not vary across time nor did time interact with the treatment effect. The combined data set was analyzed by treatment group using nominal logistic fit model (Χ^2^ = 19.7, df = 3, p < 0.001). Odds ratio analysis for treatment (Χ^2^ = 19.71, df = 3, p < 0.001) indicates that control colonies had significantly fewer queen events than fungicide (p = 0.001, CI = 4.12 - ∞) or both (p < 0.001, CI = 7.59 − ∞), but not insecticide treated (p = 0.086, CI = 0.67 - ∞) groups.

**Figure 6.**
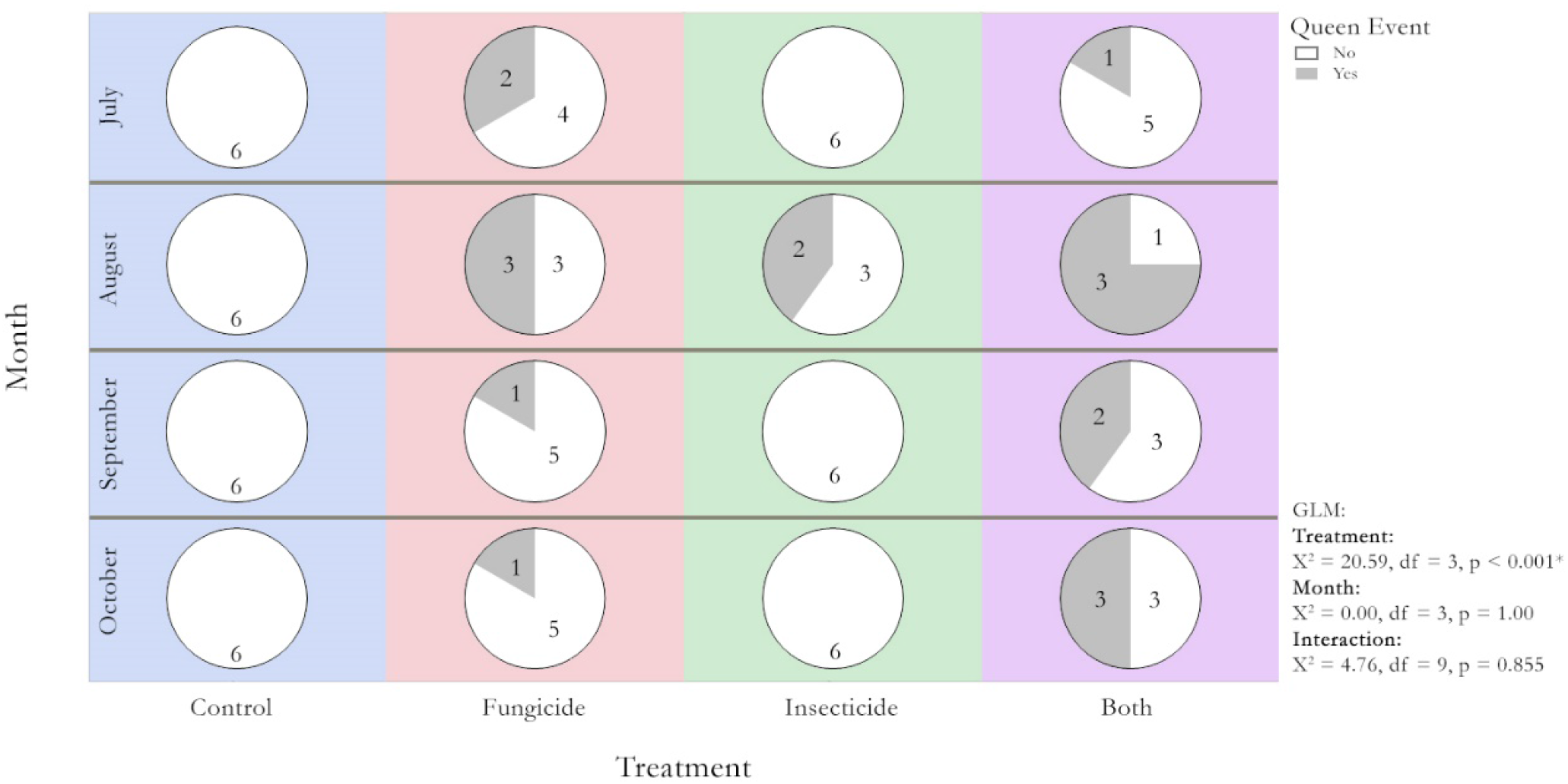
Queen Events During and After Pesticide Exposure. We exposed the colonies to sublethal pesticides in pollen for one month from July 11, 2018 until August 13, 2018. We continued to follow colony growth and queen events through October 2018, recording any queen events a colony experienced during the month. Queen events were defined as a colony lost its queen or was in the process of raising a replacement queen (queen cells with royal jelly). Colonies were managed to reduce swarm pressure, providing the colony with plenty of space for expansion. We did not intervene with any colony replacement events, allowing the colonies to requeen naturally. White = no queen event; grey = experienced a queen event. Numbers inside the circles = n colonies in each group.

## Discussion

Colonies rarely experience failure directly from pesticide exposure (21, 32), yet commercial beekeepers continue to blame pesticide exposures in crop pollination settings as the cause for increased incidences of poor brood and queen failure. National surveys of apiaries in the United States show that while pesticide contamination of pollen is widespread, the amount of pesticides detected is often at low consumption risk levels (7, 8, 43). We thus exposed colonies to field relevant contaminated pollen, dosing our pollen patties with sublethal insecticides equivalent to 10% and fungicides equivalent to 2% of a nurse bee’s lifetime LD_50_ consumption, a dose comparable to pollen contamination rates found in 15% of commercial samples (8) and 5.6% of sampled apiaries nationwide(43) and thus similar to what bees encounter in the real world.

While fungicides have been deemed bee safe as they have limited impact on adult bees and larvae, colonies exposed to them are at increased risk of experiencing queen events and long term exposure may cause atrophy of hypopharyngeal glands in a small proportion of nurse bees, which may be why we see a non-significant trend toward increased brood loss in fungicide exposed colonies (8, 43). In bumble bee colonies, exposure has resulted in smaller queens and fewer workers reared (49). The effect of fungicides on honey bee queen development is not straightforward: while fungicides in the presence of insecticides reduced queen emergence, fungicides in combination with spray adjuvants did not (50, 51). Here we demonstrate that colonies exposed to fungicides alone or in combination with insecticides experienced significantly more queen events. Why colonies exposed to sublethal fungicides engaged in increased queen replacement is still unknown and suggests a closer reexamination of fungicide exposure on queen turnover is warranted. The number of individuals with completely atrophied hypopharyngeal glands (8.7%) was small, but this extreme condition is concerning and deserves further investigation to understand the potential long-term impacts of fungicides on glands critical to nursing, glucose oxidase production critical for honey ripening (52, 53), and colony health (54, 55). Prior research has found the the herbicide glyphosate (56) as well as the fungicide pyraclostrobin, especially in conjunction with the insecticide fipronil(57) change the ultrastructure of these important glands.

Loss of larvae and reductions in brood have been previously documented when bees lack access to protein rich pollen (58–60). Colonies previously exposed to 28 days of sublethal levels of insecticides lost more brood during the larval stage, even though this brood was laid as eggs 5 days after treatment. We monitored brood during the 2^nd^ to 3^rd^ larval instar, 8 days post egg-laying, so the brood loss occurred in the second week post treatment cessation. Chlorpyrifos is known to reduce survival of larvae reared *in vitro* (61, 62). Here, and for the first time, we show that colonies exposed to pesticide products can manifest continued symptoms of that exposure into the subsequent generation of bees raised well after the exposure has stopped. Pesticide exposure to the fungicide pyraclostrobin and the insecticide fipronil has been shown to significantly impact the head proteome of nurse bees, reducing their production of royal jelly proteins, (63) which suggests pesticide exposure can reduce brood food quality and quantity. While we did not note a reduction in HPG acini size, our experiment did not investigate changes in proteome head profiles of our nurse bees across treatment groups. Alternatively, although the insecticide treated group did not consume less pollen patties than the control group, they did have a significantly higher proportion of pollen foragers, suggesting perhaps that the pesticide contaminated pollen resulted in colony wide protein stress, which could force the bees to cannibalize brood (58).

Although we did not see differences in colony strength or amount of total brood between treatment colonies, the insecticide colonies showed signs of nutritional stress. The sublethal exposure in the prior month resulted in significantly less brood reared to adulthood in the insecticide treated group, suggesting that even after colonies are removed from regions with higher pesticide exposure, such as pollination contracts (8, 64, 65), the colony can experience long-term impacts on brood viability.

Our results demonstrate that exposure to sublethal, field relevant doses of insecticides and fungicides have significant long-term effects on both queen longevity and brood viability, potentially compromising a colony’s ability to survive the winter (26). Additional research is much needed to determine why sublethal exposure to pesticides has carry-over effects on the next generation of brood rearing. In light of a growing body of research, we advocate for increased study of how sublethal insecticide exposure may cause nutritional stress, a re-evaluation of fungicides on long-term colony health and a reduction of fungicide applications during bloom times to reduce honey bee exposure.

## Materials and Methods

### Bee Colonies

We established 32 colonies from 3-lb packages in April, 2018 and built them up via feeding until colonies had expanded into two medium 1-frame Langstroth boxes at the University of Maryland Agricultural Research Station in Keedysville, Maryland. In July, 2018 we then selected 24 full-size colonies and matched them for colony strength, so that they each had similar amounts of brood and food stores. Colonies were managed using standard beekeeping practices, so that they had ample space and would not feel crowded, as overcrowding can lead to an increased swarm drive.

### Treatment Groups

The 24 matched colonies were then randomly assigned to one of four treatment groups: control, fungicide, insecticide and both. Each colony was fed four pollen patties of 80g each (320 g total). These were replaced with new patties 2x per week for four weeks. The control colonies received normal pollen patties. The fungicide colonies received pollen patties contaminated with two fungicides commonly used during blueberry pollination (chlorothalonil and propicanizole) at a sublethal contamination rate that provided the bees with an additive Hazard Quotient score (HQ) of ~200, equivalent to 2% of the honey bees LD_50_ if consumed by nurse bees for 10 days without any detoxification (8, 66). Each of the two fungicides contributed ~100 HQ. The insecticide colonies received pollen patties contaminated with the two insecticides found most commonly in commercial honey bee colonies (8) (chlorypyrifos, an organophosphate and fenpropathrin, a pyrethroid), at a sublethal contamination rate that provided the bees with an HQ score of ~1,000 (10% of LD_50_), with each insecticide contributing ~500 HQ each (Table 1 & 2). To see the specific amounts of pesticides mixed into 1 kg of pollen, see Table 2.

**Table 2.** Contamination rates of pollen patties.

| Treatment | Chemical | Tradename | LD<sub>50</sub> | HQ Contributed | Desired ppb dose | % active ingredient | Solution (in ppb) to add to 1 kg of patty | Mean dose in ppb analyzed samples sent to USDA Gastonia |
| --- | --- | --- | --- | --- | --- | --- | --- | --- |
| Fungicide | Chlorothalonil | Echo720 | 111 | 100 | 11,100 | 54.00% | 20,556 | 47,225 |
| Fungicide | Propicanizole | Fitness | 67.5 | 100 | 6,750 | 41.80% | 16,148 | 4,317 |
| **Total** |  |  |  | **200** |  |  |  | 425 + 64<br>= 489 HQ points |
| Insecticide | Chlorpyrifos | Warhawk | 0.0762 | 500 | 38.1 | 44.90% | 85 | 35 |
| Insecticide | Fenpropathrin | Danitol | 0.05 | 500 | 25 | 30.90% | 81 | 25 |
| **Total** |  |  |  | **1000** |  |  |  | 459 + 500<br>= 959 HQ points |
| Both | Chlorothalonil | Echo720 | 111 | 100 | 11,100 | 54.00% | 20,556 | 26,600 |
| Both | Propicanizole | Fitness | 67.5 | 100 | 6,750 | 41.80% | 16,148 | 5,020 |
| Both | Chlorpyrifos | Warhawk | 0.0762 | 500 | 38.1 | 44.90% | 85 | 37 |
| Both | Fenpropathrin | Danitol | 0.05 | 500 | 25 | 30.90% | 81 | 25 |
| **Total** |  |  |  | **1,200** |  |  |  | 239 + 74 + 486 + 500<br>= 1299 HQ point |

\* We analyzed several test samples of pollen (n = 7) at the USDA Gastonia lab for pesticide contamination using the off-the-shelf plant protection chemicals farmers apply to fields. The chlorothalonil detected in our samples was significantly higher than expected, while the other pesticides we incorporated were close to our intended dose. All pesticides were purchased in the formulation products available to farmers from Southern States.

### Pollen Consumption

At the beginning of each inspection (every 7 days, unless delayed by rain) - any remainder of the pollen patties was collected into individual Ziploc bags and replaced with fresh patties. The remainder was weighed and subtracted from 320 g to determine consumption.

### Pollen Trapping

Each colony was fitted with a Sundance Pollen Trap (Ross Rounds, Canandaigua, NY) to minimize the amount of fresh, potentially uncontaminated pollen a colony could collect. These traps were constantly engaged over the first 4 weeks of the study, during which collonies were provisioned with pollen patties. This helped ensure that colonies were dependent on the pollen patties for protein. We collected pollen from these traps on five separate occasions during the month we fed contaminated pollen patties. Five days after we ceased feeding contaminated pollen, the traps were opened so that the bees could forage freely and bring back fresh pollen.

### Colony Foraging

The entrances were reduced with wire mesh for 5-minute intervals on four separate days. An observer watched the entrance and recorded all returning foragers on two hand counters, one for pollen foragers and one for non-pollen foragers (39, 46).

### Brood Viability

Once we removed the final contaminated pollen patties, we caged each queen on an empty comb for 24 hours to produce eggs of a specific age. We then followed the viability of 50 eggs through development until adult emergence. These larvae are fed by nurse bees. Bees typically engage in nursing between 5 and 14 days post emergence as an adult (12, 67). Worker bee development from egg to adult takes 21 days, so at the time we ceased feeding the contaminated pollen patties after 28 days, all of the nurse bees that were of an appropriate nursing age had been reared in a colony experiencing sublethal pesticide exposure during their development and hence we were measuring the carry-over effects of pesticide exposure on the next generation reared.

### Hypopharyngeal Glands

The bees that emerged from the brood viability test were marked with paint on the thorax and allowed to age in their respective colonies. They were recaptured at 7 days of age for hypopharyngeal dissections, stored at −80 °C until dissected. The glands were dissected into a concave slide and the width of glands measured using an Amscope 10 megapixel color camera, model MU1000 and processed with Amscope 3.7 digital software, using the protocol outlined in Renzi (48).

### Colony Size and Queen Events

The queen is the sole reproductive individual in a colony and her pheromones regulate colony cohesion. We marked all of our queens at the start of the experiment. Colonies were evaluated on a monthly basis for the number of brood frames, the number of frames occupied by bees, and if the colonies experienced a queen event, indicated by queenlessness or the presence of a queen cell containing a royal jelly provisioned queen cell. Sometimes colonies will rear a supersedure queen. During this supersedure process there can be two queens in the same colony and then the original one is eventually killed off; no successful supercedure events occurred during this trial.

### Statistics

All statistical analysis were conducted using JMP Pro 14.1.0. We conducted a generalized linear model with appropriate distribution for pollen consumption, colony wide pollen collection, number of foragers, individual brood stages, and queen events. Since treatment and date did not interact for number of foragers, we pooled all data by treatment group to compare the number of pollen foragers vs. non-foragers in a 2×4 contingency table. We analyzed brood loss over time using a repeated measures MANOVA, we found a significant effect of both time and treatment, and no interaction. As there was no interaction with time, we analyzed each stage of development separately, comparing loss rates across treatment groups using a generalized linear model with normal distribution. Since month was not significant for queen events, we pooled queen events for the entire season and analyzed by treatment group using nominal logistic fit. As this was significant, we conducted an odds ratio analysis for each group compared to controls.

## Acknowledgements

A NIFA ELI postdoctoral fellowship (grant no. 2015-03505) from the USDA National Institute of Food and Agriculture and NE SARE project LNE19-392R-3324 supported this work.

## Notes

### Competing Interest Statement

The authors have declared no competing interest.

## References

1. J. Ollerton, R. Winfree, S. Tarrant, How many flowering plants are pollinated by animals? Oikos 120, 321–326 (2011).

2. A.-M. Klein et al., Importance of pollinators in changing landscapes for world crops. Proceedings of the royal society B: biological sciences 274, 303–313 (2007).

3. N. Gallai, J.-M. Salles, J. Settele, B. E. Vaissière, Economic valuation of the vulnerability of world agriculture confronted with pollinator decline. Ecological Economics 68, 810–821 (2009).

4. N. W. Calderone, Insect Pollinated Crops, Insect Pollinators and US Agriculture: Trend Analysis of Aggregate Data for the Period 1992–2009. PloS ONE 7, e37235 (2012).

5. D. Atwood, C. Paisley-Jones (2017) Pesticides Industry Sales and Usage, 2008 – 2012, Market Estimates. ed E. P. Agency (Washington, DC).

6. B. Chèze, M. David, V. Martinet, Understanding farmers’ reluctance to reduce pesticide use: A choice experiment. Ecological Economics 167, 106349 (2020).

7. C. A. Mullin et al. High Levels of Miticides and Agrochemicals in North American Apiaries: Implications for Honey Bee Health. PLoS ONE 5, e9754 (2010).

8. K. S. Traynor et al. In-hive Pesticide Exposome: Assessing risks to migratory honey bees from in-hive pesticide contamination in the Eastern United States. Scientific reports 6, 33207 (2016).

9. E. Carnesecchi et al., Investigating combined toxicity of binary mixtures in bees: Meta-analysis of laboratory tests, modelling, mechanistic basis and implications for risk assessment. Environment International 133, 105256 (2019).

10. K. Crailsheim, The flow of jelly within a honeybee colony. Journal of Comparative Physiology B 162, 681–689 (1992).

11. T. Schmickl, K. Crailsheim, Inner nest homeostasis in a changing environment with special emphasis on honey bee brood nursing and pollen supply. Apidologie 35, 249–263 (2004).

12. K. Crailsheim et al., Pollen consumption and utilization in worker honeybees (Apis mellifera carnica): Dependence on individual age and function. Journal of Insect Physiology 38, 409–419 (1992).

13. Anonymous (2012) White Paper in Support of the Proposed Risk Assessment Process for Bees. (Office of Chemical Safety and Pollution Prevention, Washington, DC), p 275.

14. F. Böhme, G. Bischoff, C. P. W. Zebitz, P. Rosenkranz, K. Wallner, From field to food—will pesticide-contaminated pollen diet lead to a contamination of royal jelly? Apidologie 49, 112–119 (2018).

15. D. Goulson, E. Nicholls, C. Botías, E. L. Rotheray, Bee declines driven by combined stress from parasites, pesticides, and lack of flowers. Science 347, 1255957 (2015).

16. F. Sanchez-Bayo, K. Goka, Pesticide residues and bees–a risk assessment. PloS one 9, e94482 (2014).

17. V. Doublet, M. Labarussias, J. R. de Miranda, R. F. Moritz, R. J. Paxton, Bees under stress: sublethal doses of a neonicotinoid pesticide and pathogens interact to elevate honey bee mortality across the life cycle. Environmental microbiology 17, 969–983 (2015).

18. C. Alaux et al., Interactions between *Nosema* microspores and a neonicotinoid weaken honeybees (*Apis mellifera*). Environmental microbiology 12, 774–782 (2010).

19. C. Vidau et al., Exposure to Sublethal Doses of Fipronil and Thiacloprid Highly Increases Mortality of Honeybees Previously Infected by Nosema ceranae. PLoS ONE 6, e21550 (2011).

20. J. S. Pettis et al. Crop Pollination Exposes Honey Bees to Pesticides Which Alters Their Susceptibility to the Gut Pathogen Nosema ceranae. PLOS ONE 8, e70182 (2013).

21. F. Sánchez-Bayo et al., Are bee diseases linked to pesticides?—A brief review. Environment international 89, 7–11 (2016).

22. N. Simon-Delso et al., Honeybee colony disorder in crop areas: the role of pesticides and viruses. PloS one 9, e103073 (2014).

23. Y. Poquet, C. Vidau, C. Alaux, Modulation of pesticide response in honeybees. Apidologie 47, 412–426 (2016).

24. D. R. Schmehl, P. E. Teal, J. L. Frazier, C. M. Grozinger, Genomic analysis of the interaction between pesticide exposure and nutrition in honey bees (Apis mellifera). Journal of insect physiology 71, 177–190 (2014).

25. S. Tosi, J. C. Nieh, F. Sgolastra, R. Cabbri, P. Medrzycki, Neonicotinoid pesticides and nutritional stress synergistically reduce survival in honey bees. Proceedings of the Royal Society B: Biological Sciences 284, 20171711 (2017).

26. D. vanEngelsdorp, D. R. Tarpy, E. J. Lengerich, J. S. Pettis, Idiopathic brood disease syndrome and queen events as precursors of colony mortality in migratory beekeeping operations in the eastern United States. Preventive Veterinary Medicine 108, 225–233 (2013).

27. S. T. O’Neal, T. D. Anderson, J. Y. Wu-Smart, Interactions between pesticides and pathogen susceptibility in honey bees. Current Opinion in Insect Science 26, 57–62 (2018).

28. M. Henry et al., A Common Pesticide Decreases Foraging Success and Survival in Honey Bees. Science 336, 348–350 (2012).

29. M. A. Becher, J. L. Osborne, P. Thorbek, P. J. Kennedy, V. Grimm, REVIEW: Towards a systems approach for understanding honeybee decline: a stocktaking and synthesis of existing models. Journal of Applied Ecology 50, 868–880 (2013).

30. N. Steinhauer et al., Drivers of colony losses. Current Opinion in Insect Science 26, 142–148 (2018).

31. K. Kulhanek et al., A national survey of managed honey bee 2015–2016 annual colony losses in the USA. Journal of Apicultural Research 56, 328–340 (2017).

32. G. P. Dively, M. S. Embrey, A. Kamel, D. J. Hawthorne, J. S. Pettis, Assessment of chronic sublethal effects of imidacloprid on honey bee colony health. PloS one 10(2015).

33. P. Calatayud-Vernich, F. Calatayud, E. Simó, M. M. Suarez-Varela, Y. Picó, Influence of pesticide use in fruit orchards during blooming on honeybee mortality in 4 experimental apiaries. Science of the Total Environment 541, 33–41 (2016).

34. O. Samson-Robert, G. Labrie, M. Chagnon, V. Fournier, Planting of neonicotinoid-coated corn raises honey bee mortality and sets back colony development. PeerJ 5(2017).

35. L. Straub, G. R. Williams, J. Pettis, I. Fries, P. Neumann, Superorganism resilience: eusociality and susceptibility of ecosystem service providing insects to stressors. Current Opinion in Insect Science 12, 109–112 (2015).

36. R. Brodschneider, R. Moosbeckhofer, K. Crailsheim, Surveys as a tool to record winter losses of honey bee colonies: a two year case study in Austria and South Tyrol. Journal of Apicultural Research 49, 23–30 (2010).

37. A. M. Spleen et al. A national survey of managed honey bee 2011–12 winter colony losses in the United States: results from the Bee Informed Partnership. Journal of Apicultural Research 52, 44–53 (2013).

38. T. Schuler (2018) Pollination in blueberries.

39. K. S. Traynor, Y. Le Conte, R. E. Page Jr, Age matters: pheromone profiles of larvae differentially influence foraging behaviour in the honeybee, Apis mellifera. Animal behaviour 99, 1–8 (2015).

40. J. Free, A. Ferguson, J. R. Simpkins, Influence of virgin queen honeybees (Apis mellifera) on queen rearing and foraging. Physiological entomology 10, 271–274 (1985).

41. R. Rader et al., Alternative pollinator taxa are equally efficient but not as effective as the honeybee in a mass flowering crop. Journal of Applied Ecology 46, 1080–1087 (2009).

42. A. Sáez, C. L. Morales, L. Y. Ramos, M. A. Aizen, Extremely frequent bee visits increase pollen deposition but reduce drupelet set in raspberry. Journal of Applied Ecology 51, 1603–1612 (2014).

43. K. Traynor et al., Pesticides in Honey Bee Colonies: real world exposure and associated morbidity over seven years (2011-2017) in the USA. ((submitted)).

44. United States Geological Service (2020) Estimated Annual Agricultural Pesticide Use: Pesticide Use Maps. ed National Water-Quality Assessment (NAWQA) Project.

45. K. S. Delaplane, J. J. M. van der Steen, E. Guzman-Novoa, K. S. Delaplane, Standard methods for estimating strength parameters of *Apis mellifera* colonies. Journal of Apicultural Research 52, 1–12 (2013).

46. K. S. Delaplane et al. Standard methods for pollination research with Apis mellifera. Journal of Apicultural Research 52, 1–28 (2013).

47. C. Dreller, R. E. Page Jr, M. K. Fondrk, Regulation of pollen foraging in honeybee colonies: effects of young brood, stored pollen, and empty space. Behavioral Ecology and Sociobiology 45, 227–233 (1999).

48. M. T. Renzi et al. Combined effect of pollen quality and thiamethoxam on hypopharyngeal gland development and protein content in Apis mellifera. Apidologie 47, 779–788 (2016).

49. O. M. Bernauer, H. R. Gaines-Day, S. A. Steffan, Colonies of bumble bees (Bombus impatiens) produce fewer workers, less bee biomass, and have smaller mother queens following fungicide exposure. Insects 6, 478–488 (2015).

50. G. DeGrandi-Hoffman, Y. Chen, R. Simonds, The effects of pesticides on queen rearing and virus titers in honey bees (Apis mellifera L.). Insects 4, 71–89 (2013).

51. R. M. Johnson, E. G. Percel, Effect of a fungicide and spray adjuvant on queen-rearing success in honey bees (Hymenoptera: Apidae). Journal of economic entomology 106, 1952–1957 (2013).

52. T. Takenaka, H. Ito, K. Yatsunami, T. Echigo, Changes of glucose oxidase activity and amount of gluconic acid formation in the hypopharyngeal glands during the lifespan of honey bee workers (Apis mellifera L.). Agricultural and biological chemistry 54, 2133–2134 (1990).

53. K. Ohashi, S. Natori, T. Kubo, Expression of amylase and glucose oxidase in the hypopharyngeal gland with an age-dependent role change of the worker honeybee (Apis mellifera L.). European Journal of Biochemistry 265, 127–133 (1999).

54. D.-I. Wang, F. Moeller, Histological comparisons of the development of hypopharyngeal glands in healthy and Nosema-infected worker honey bees. Journal of Invertebrate Pathology 14, 135–142 (1969).

55. S.-C. Seehuus, K. Norberg, T. Krekling, K. Fondrk, G. V. Amdam, Immunogold localization of vitellogenin in the ovaries, hypopharyngeal glands and head fat bodies of honeybee workers, *Apis mellifera*. Journal of Insect Science 7, 52 (2007).

56. M. R. Faita, E. de Medeiros Oliveira, V. V. A. Júnior, A. I. Orth, R. O. Nodari, Changes in hypopharyngeal glands of nurse bees (Apis mellifera) induced by pollen-containing sublethal doses of the herbicide Roundup®. Chemosphere 211, 566–572 (2018).

57. R. Zaluski, L. A. Justulin, R. de Oliveira Orsi, Field-relevant doses of the systemic insecticide fipronil and fungicide pyraclostrobin impair mandibular and hypopharyngeal glands in nurse honeybees (Apis mellifera). Scientific reports 7, 1–10 (2017).

58. T. Schmickl, K. Crailsheim, Cannibalism and early capping: strategy of honeybee colonies in times of experimental pollen shortages. Journal of Comparative Physiology A 187, 541–547 (2001).

59. H. N. Scofield, H. R. Mattila, Honey bee workers that are pollen stressed as larvae become poor foragers and waggle dancers as adults. Plos one 10, e0121731 (2015).

60. F. Requier, J. F. Odoux, M. Henry, V. Bretagnolle, The carry-over effects of pollen shortage decrease the survival of honeybee colonies in farmlands. Journal of applied ecology 54, 1161–1170 (2017).

61. P. Dai, C. J. Jack, A. N. Mortensen, J. D. Ellis, Acute toxicity of five pesticides to Apis mellifera larvae reared in vitro. Pest management science 73, 2282–2286 (2017).

62. P. Dai et al., Chronic toxicity of clothianidin, imidacloprid, chlorpyrifos, and dimethoate to Apis mellifera L. larvae reared in vitro. Pest management science 75, 29–36 (2019).

63. Z. Rodrigo et al., Modification of the head proteome of nurse honeybees (Apis mellifera) exposed to field-relevant doses of pesticides. Scientific Reports (Nature Publisher Group) 10(2020).

64. N. Carreck, P. Neumann, Honey bee colony losses. J Apic Res 49, 1–6 (2010).

65. F. Böhme, G. Bischoff, C. P. W. Zebitz, P. Rosenkranz, K. Wallner, Pesticide residue survey of pollen loads collected by honeybees (Apis mellifera) in daily intervals at three agricultural sites in South Germany. PLOS ONE 13, e0199995 (2018).

66. K. A. Stoner, B. D. Eitzer, Using a hazard quotient to evaluate pesticide residues detected in pollen trapped from honey bees (Apis mellifera) in Connecticut. 8, e77550 (2013).

67. G. A. Rösch, Untersuchungen über die Arbeitsteilung im Bienenstaat. Zeitschrift für vergleichende Physiologie 2, 571–631 (1925).

