## Supplemental Figures 1-3 for "Social Disruption: Sublethal pesticides in pollen lead to *Apis mellifera* queen events and brood loss"

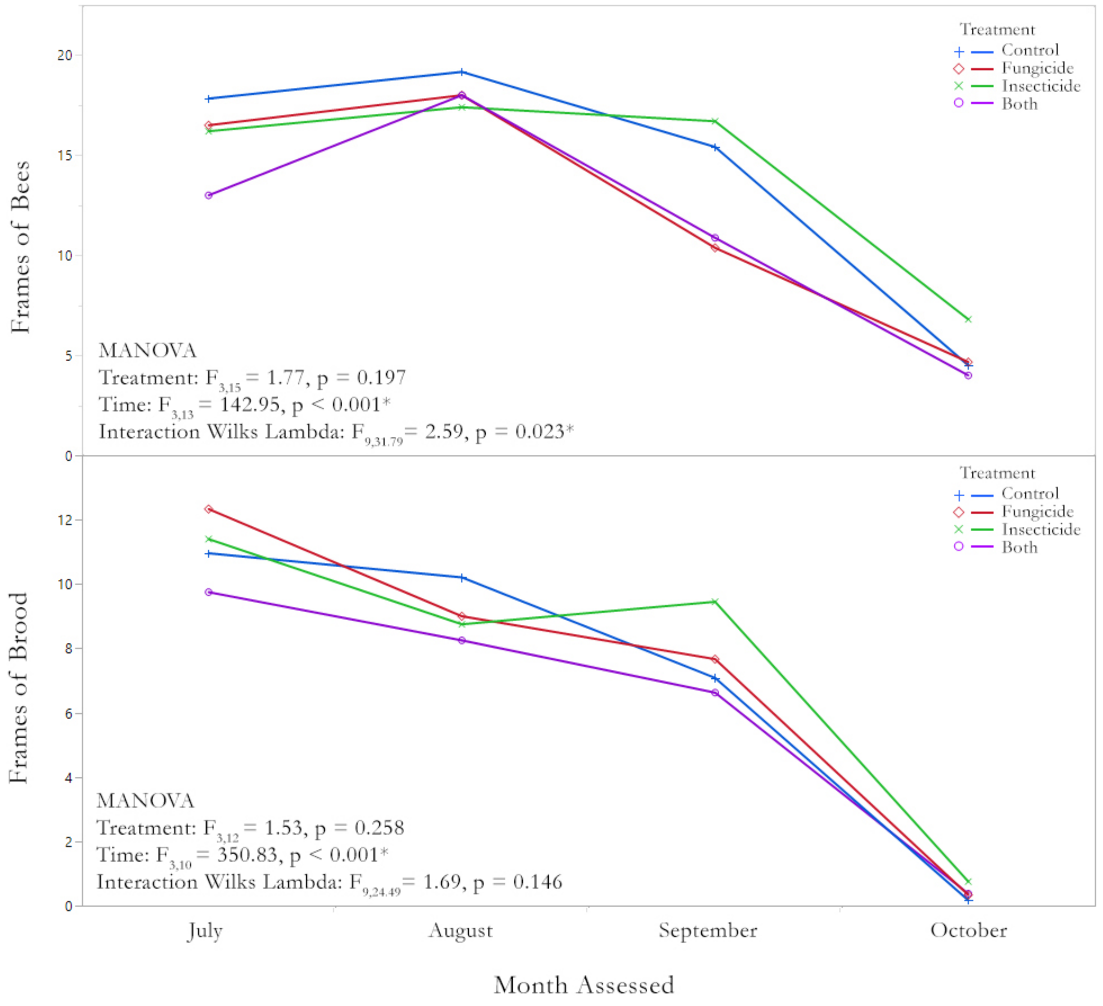


**Supplemental Figure 1.** Frames of Bees and Brood over Time

Repeated measures analysis of frames of bees and frames of brood analyzed over time. For both bees and brood there was a significant effect of time, but no effect of treatment. There was an interaction effect of treatment and time for frames of bees, but not for frames of brood. Control = blue, fungicide = red, insecticide = green, both = purple.

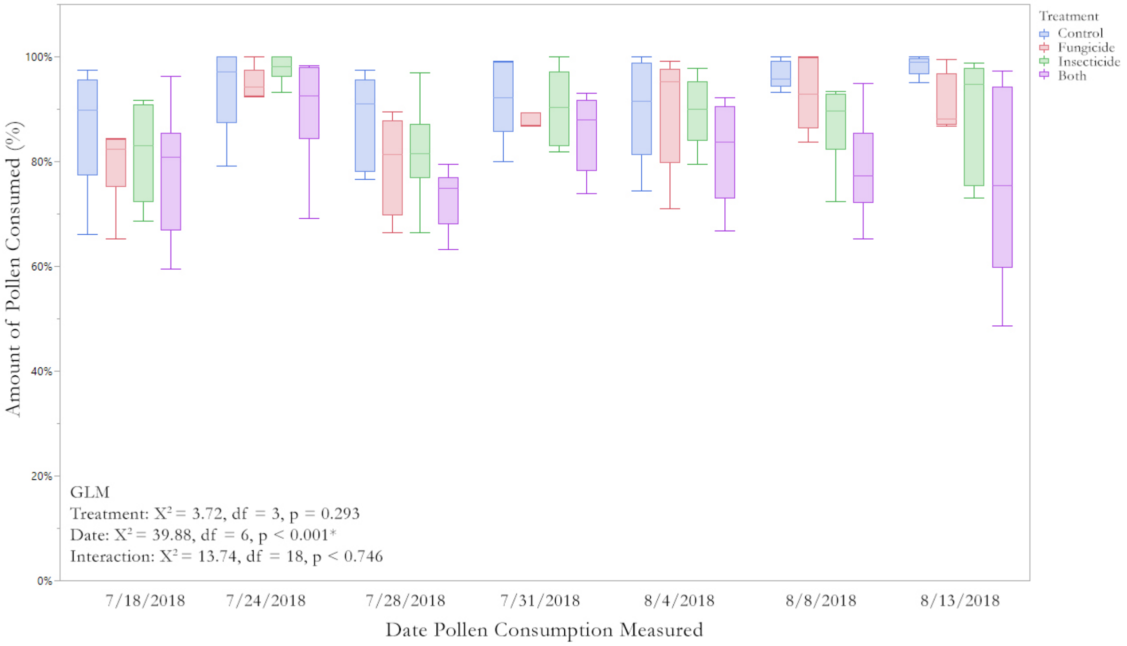


**Supplemental Figure 2.** Pollen Consumption by Date and Treatment Group

Each colony was provided with four 80g fresh pollen patties 2x per week, typically on the Tues and Saturday, unless it rained. At each indicated date the remainder of the patties was removed and replaced with fresh ones. Colonies could thus consume up to 320g of pollen per feeding. Consumption was analyzed using a GLM with treatment and date as factors. Pollen consumption differed over time, but did not vary significantly by treatment group. Control = blue, fungicide = red, insecticide = green, both = purple. Box plots show quartiles and outliers.

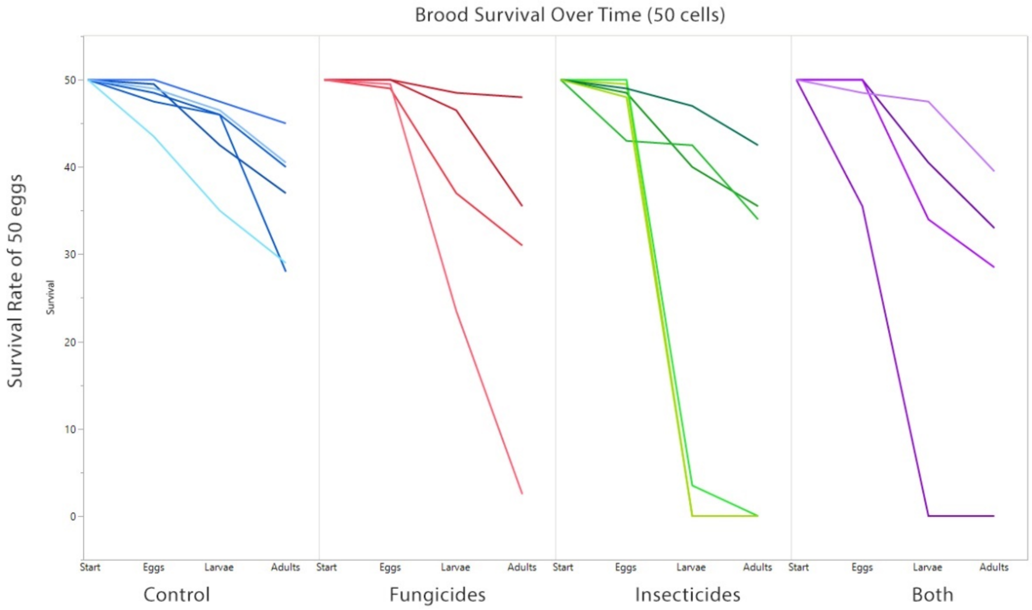


**Supplemental Figure 3.** Brood Loss by Individual Colony

Here we show the brood loss and/or cannibalism for each individual colony, segregated by treatment group. All of the colonies in the control group had at least 50% their brood survive, with the majority losing less than 10 cells during the 21 days of development, whereas all three treatment groups had at least one colony that removed/cannibalized all 50 cells and had several where at least 15 cells were lost. Only in the insecticide and both group did we see total brood loss in the larval stage. Control = blue, fungicide = red, insecticide = green, both = purple.
